# Identification of Freezing Tolerance QTLs in *Tripsacum dactyloides* Using Open-Pollinated Bulk Segregant Analysis

**DOI:** 10.64898/2025.12.10.693030

**Authors:** Mohamed Z. El-Walid, Christine M. Gault, Denise E. Costich, Nicholas K. Lepak, Joshua S. Budka, Michelle C. Stitzer, Anju Giri, Evan Rees, M Cinta Romay, Edward S. Buckler, Sheng-Kai Hsu

**Author notes:** Corresponding Authors: Mohamed Z. El-Walid, Edward S. Buckler, Sheng-Kai Hsu.

## Abstract

This study investigates the genetic basis of freezing tolerance in *Tripsacum dactyloides* and related subspecies as a potential source of valuable traits for improving maize agriculture. Recognizing the significant economic losses in corn yields due to frost damage, we hypothesized that northern populations of *T. dactyloides* are enriched for freezing tolerance alleles. 40 diverse *Tripsacum* accessions were collected from natural populations and long-established field collections and used to generate F1 hybrids and open-pollinated F2 families. F2 seedlings were germinated then screened within a growth chamber for freezing tolerance by exposure to freezing temperatures. Seedlings were then phenotyped by tissue survival, and extremes were pooled to create tolerant and susceptible bulks. DNA sequencing was performed on founders, F1s, and tolerant/susceptible F2 bulks. To overcome challenges in traditional SNP calling in bulked samples, we developed a regression-based approach to estimate gamete frequencies and impute allele frequencies in pooled populations. The results showed genetic diversity among Tripsacum accessions, with divergence between northern and southern populations. We tracked segregation of alleles across genomic loci, and performed a joint bulk segregant analysis, identifying 9 QTLs significantly associated with freezing tolerance. These findings highlight potential loci for freezing tolerance that could inform genetic engineering of maize.

**Central Hypothesis:** Northern populations of *Tripsacum dactyloides*, a wild relative of maize, are enriched for freezing tolerance alleles which can be identified by mapping.

## Introduction

### Freezing Tolerance May Substantially Reduce Corn Losses

Freezing tolerance in corn could significantly benefit agriculture by enabling earlier planting and extending the growing season. Planting earlier helps crops avoid high temperatures during flowering, while a longer season allows for earlier canopy closure, increased solar radiation capture, and greater biomass accumulation, as seen in the higher biomass yield of Miscanthus compared to maize [1]. Furthermore, frost tolerance may lead to more efficient fertilizer use due to quicker crop establishment and provide a competitive edge against weeds. Overall, incorporating frost tolerance in corn could lead to more dependable yields through enhanced planting flexibility and reduced exposure to heat stress during critical growth stages, alongside improved agronomic efficiency via decreased input requirements.

In 2024, the United States allocated about one-third of its total crop acreage to corn, totaling approximately 91 million acres. This corn cultivation is projected to yield grains valued at around $64.7 billion [2]. While quantifying the yearly economic impact of abiotic stresses is difficult, research indicates that early-season light frost can cause up to 8% in corn grain yield reductions, and extended drought can lead to a daily yield loss exceeding 9% [3,4]. Cumulatively, these abiotic stresses can result in tens of billions of dollars in lost grain yield annually. Therefore, even slight improvements in mitigating their effects could generate billions of dollars in additional yearly crop yields. The creation of freeze-tolerant corn offers a promising avenue to achieve this goal.

### Plant physiological response to cold

In terrestrial plants, exposure to subzero temperatures imposes significant physiological stress. This stress manifests primarily through two avenues: water deprivation resulting from extracellular ice formation in the soil and on the plant’s surface, and the disruption of cellular integrity and function due to intracellular freezing, which can lead to membrane perforation, detrimental phase changes in lipids, and protein denaturation [5,6].

Plants have broadly evolved three strategies to contend with freezing conditions: temporal avoidance through phenological shifts, sacrificing exposed tissues followed by regrowth from insulated structures like rhizomes, and the development of molecular mechanisms for tolerance [7–9]. It is this third strategy, the ability to withstand freezing at a cellular level, that holds particular relevance for enhancing the viability of early-season maize plantings that may face acute, overnight freezing events.

### Tripsacum dactyloides is the most promising source of transferable freezing tolerance

While robust freezing tolerance is lacking in existing maize diversity, it has evolved in related grass species. *Tripsacum dactyloides* is a perennial native to the North American grasslands, and spans climates from Mexico and Florida to New York and Iowa. Observations in Tripsacum reveal accelerated evolution in stress-response genes, particularly within phospholipid metabolism pathways, which are critical for maintaining membrane fluidity at low temperatures [10]. *Tripsacum dactyloides* is also a close relative of maize, having diverged only 650 thousand years ago and share their most recent whole genome duplication event [11,12].

We observed minimal cold tolerance in *Tripsacum dactyloides* subsp. *floridanum*, a taxon geographically restricted to the southern tip of Florida [13], with probable origins as an isolated tropical Pleistocene refugia population [14]. We hypothesize that northern populations of *Tripsacum dactyloides* are, in contrast, enriched for genetically mappable freezing tolerance alleles. For these reasons, *Tripascum dactyloides* is a good candidate to source freezing tolerance that is transferable to maize.

## Methods

### Hypothesis and Experimental Design

Our goal in this study is to identify the genetic architecture of seedling freezing tolerance. In order to meet this goal, we generate a mapping population, screen for freezing tolerance, and establish a bulk segregant genetic mapping strategy for an understudied outcrossing wild species (Figure 1). This strategy integrates both linkage (recent recombination) and association (ancestral recombination) approaches, similar to those used in the maize NAM design (Yu et al., 2008), but with bulk segregant analysis. Given the recent divergence between *Zea* (freezing intolerant) and *Tripsacum* (freezing tolerant), we hypothesize that relatively few large-effect freezing tolerance alleles segregate in North American *T. dactyloides* populations. We aim to identify these loci and test whether they have experienced differential selection across latitudes.

**Figure 1:**
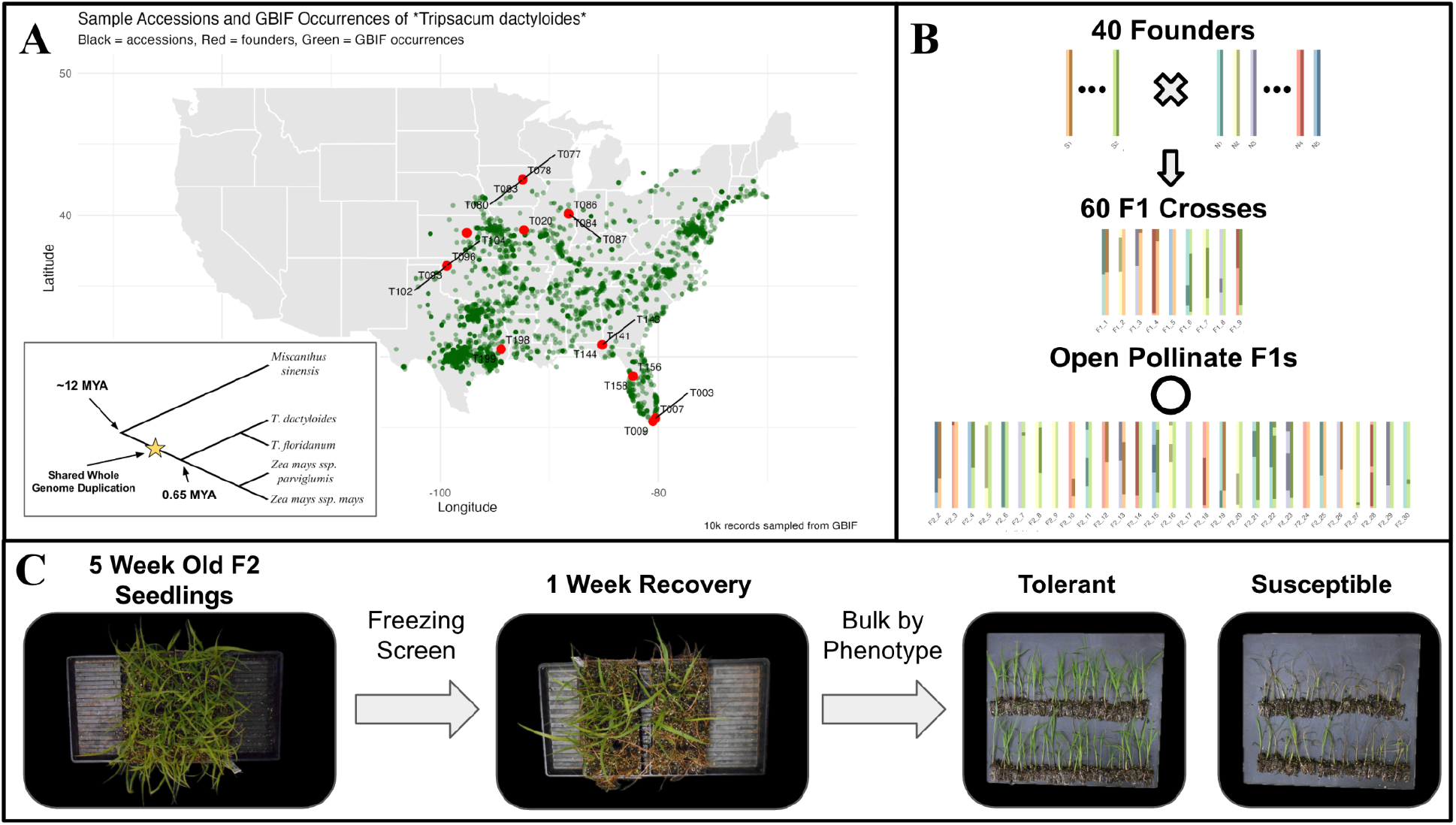
A) Depiction of the continental United States with iNaturalist research-grade observations of *Tripsacum dactyloides* shown in green and sampling locations for each mapping population founder shown in red. Includes a depiction of a phylogenetic tree showing the evolutionary relationship between *Miscanthus sinensis, Tripsacum dactyloides subsp. dactyloides, Tripsacum dactyloides subsp. floridanum*, and *Zea mays ssp. mays*. B) Cartoon depiction of the crossing scheme for the *Tripsacum* mapping population and examples of possible chromosomal lineages based on simulated recombination. C) Images showcasing *Tripsacum* seedlings before and after exposure to freezing. Also includes examples of a tolerant bulk and a susceptible bulk that were pooled and sequenced.

**Figure 2:**
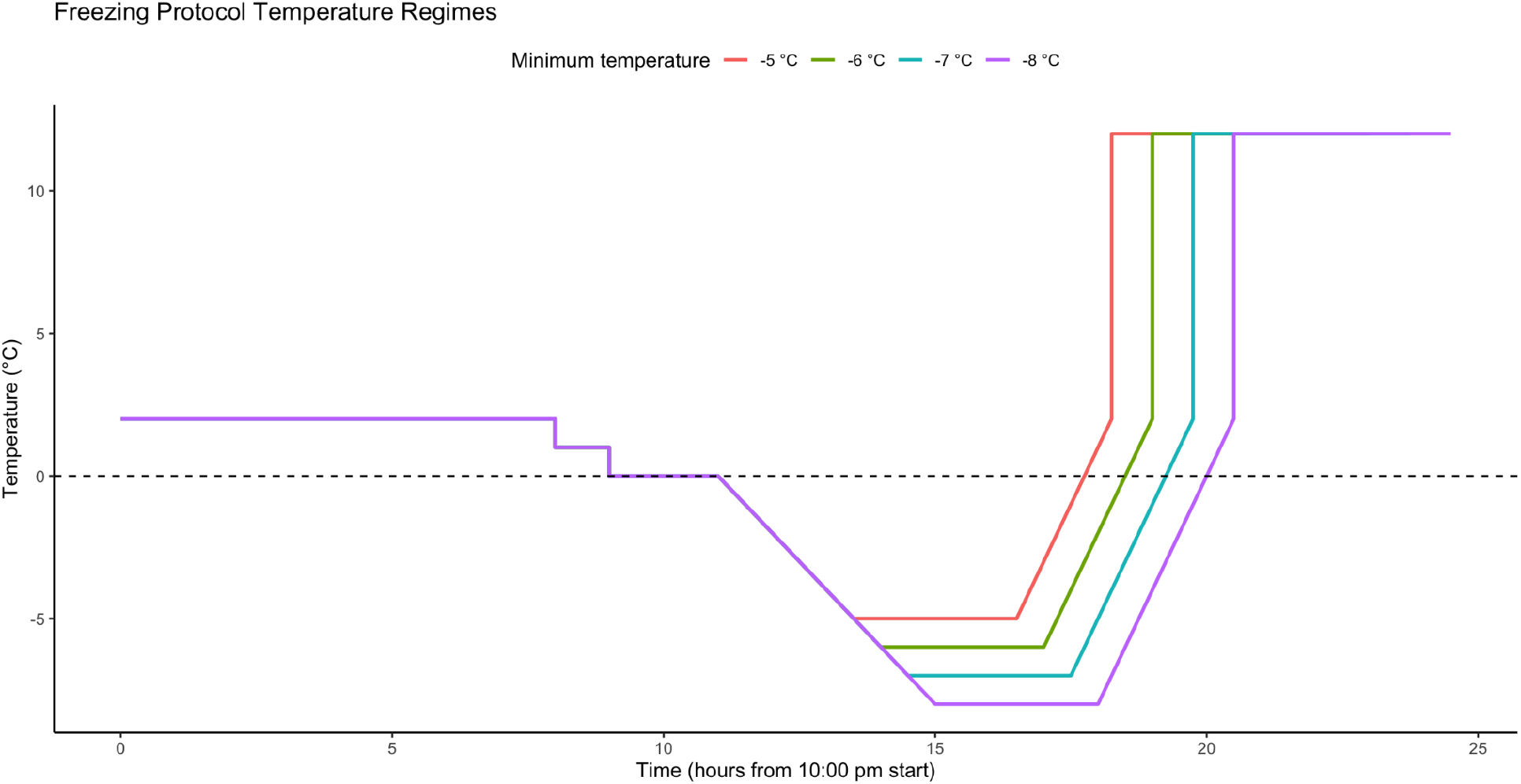
Depiction of freezing screen temperature regimes over time. The regime for each screen was selected based on the maternal parent’s freezing tolerance to improve the stratification between tolerant and susceptible seedlings.

### Germplasm, Crossing, and Seed generation

*Tripsacum dactyloides* subsp. *dactyloides* and *Tripsacum dactyloides* subsp. *floridanum* samples were collected as rhizomes from across the continental United States to serve as our population founders (Sup Table 1). The clones were chosen to maximize temperature and latitudinal ranges, with founders designated as northern if they were collected from northern latitudes that experience heavy freezing and frost conditions, and founders designated as southern if they were collected from southern latitudes with sub-tropical climates. Most clones were stored temporarily in a cold room at 4°C for several weeks. During the storage period, damage to some Florida clones was observed suggesting a difference in chilling tolerance even among the founder germplasm. Based on subspecies, ploidy, environmental background, and this informal chilling stress test, 40 parents were chosen to maximize segregation of potential frost tolerance QTL by crossing *Tripsacums* of northern origin by *Tripsacums* of southern origin.

The resulting 60 F1 crosses were clonally propagated and planted in replicates of 12 plants in a common field in Ithaca, NY (42.449973, -76.461485). All F1s were able to survive winters in Ithaca. F2 seeds produced from open-pollination of F1 individuals in the field were collected for further screening.

### Phenotypic screen

Phenotypic screens for freezing tolerance were conducted using F2 seed. Seeds were sown into 288-well trays of Cornell Mix (7.6 ft^3^ of peat, 30 lbs of vermiculite, 15 lbs of perlite, 5 lbs of lime, 4 lbs 10-5-10 fertilizer and 4 lbs calcium sulfate). Trays were placed on heating mats set to 80℉ and grown for ∼4 weeks under standard greenhouse conditions (14 hours at 28°C in the light, where supplemental lights turned on when natural light was below 500 W m-2, and 10 hours at 22°C in the dark in summer (April through October); 11 hours at 24°C in the light and 13 hours at 18°C in the dark in winter (November through March)). Seedlings were then transplanted into 128-well trays to reestablish. Five weeks after sowing, seedlings were cold-acclimated in a growth chamber (12°C, 600 umol/m2/s at 1 foot high, 13 hour photoperiod from 7am-8pm EST) for one week. After cold acclimation, sickly seedlings (potentially due to inbreeding depression) were removed and the remaining cold acclimated plants were treated with SnowMax to promote the nucleation of ice crystals. The soil was soaked in a 1x SnowMax solution and leaf tissue was also treated with 1x SnowMax solution.

After cold acclimation, seedling trays were placed in a freezer set at 2 °C overnight with the lights off, which allowed the soil to reach thermal equilibrium. Two temperature probes recorded the temperature at mid-plant height every 30 seconds. The freezer contained an electronic rotary tray to expose all seedlings equally to different microenvironments in the chamber. At 6 am the next morning, the temperature decreased to 1 °C for one hour. At 7 am, the temperature decreased to 0 °C for two hours. Just before 9 am, ice chips were spread on the soil so that every seedling stem was in contact with ice. Starting at 9 am, the lights turned on and the freezer ramped down to the minimum freezing temperature. A minimum freezing temperature was selected for each batch according to the freezing tolerance of the maternal F1 parent. When the minimum temperature was set to -5 °C, it took 2.5 hours to ramp from 0 °C to -5 °C. After a three hour exposure at -5 °C, the temperature ramped up to 2 °C over 1 hour and 45 minutes. When the minimum temperature was set to -6 °C, it took 3 hours to ramp from 0 °C to -6 °C. After a three hour exposure at -6 °C, the temperature ramped up to 2 °C over two hours. When the minimum temperature was set to -7 °C, it took 3.5 hours to ramp from 0 °C to -7 °C. After a three hour exposure at -7 °C, the temperature ramped up to 2 °C over 2 hours and 15 minutes. When the minimum temperature was set to -8 °C, it took four hours to ramp from 0 °C to -8 °C. After a three hour exposure at -8 °C, the temperature ramped up to 2 °C over 2 hours and 30 minutes. The tray was then transferred to the 12 °C cold acclimation chamber. After 24 hours at 12 °C, the tray was moved to the greenhouse and allowed to recover for six days.

Phenotyping and tissue collection were performed six days after freezing exposure. The tray was divided into eight sections of approximately 16 seedlings to further account for micro environment variability in the freezing chamber. Seedling freezing tolerance was visually scored based on leaf tissue survival, and the top and bottom 15-20% of seedlings were then classified as the freezing tolerant and susceptible bulks respectively. For each seedling in the tolerant bulk, one leaf hole-punch was collected and pooled with their respective bulk. For the freezing-susceptible bulk, an equal amount of root hair tissue was sampled from each selected seedling.

Various temperature regimes were tried for each F1 family to determine the minimum freezing temperature according to each family’s freezing tolerance, which allowed for phenotypic segregation of freezing tolerance. In rare cases, the freezing exposure was too harsh or too mild. In cases where more than a third of the seedlings were completely dead, a freezing-susceptible bulk was not collected, and only a freezing-tolerant bulk was collected. In cases where more than a third of the seedlings were perfectly healthy, a freezing-tolerant bulk was not collected, and only a freezing-susceptible bulk was collected. Tissue samples were then stored at -80℃ prior to DNA extraction.

### DNA Extraction and Sequencing

Leaf tissue was collected from mature clones of each of the founders and each of the F1s. To generate the pools, leaf tissue was collected from tolerant seedlings, and root tissue from susceptible seedlings post screening. DNA isolation was done using a Qiagen DNeasy Plant DNA extraction kit. DNA was sequenced using a combination of HiSeq X Ten, HiSeq 4000, and Illumina NovaSeq 6000 through Novogene. Sequencing depth and number of libraries varied (S Table 5: Mean sequencing depth by sample).

107 bulks containing 3-66 pooled seedlings were skim sequenced at 1-2X coverage, on HiSeq X Ten or HiSeq 4000 through Novogene (Sup Table 3).

### Sequence Alignment

We use two high-quality, haplotype-resolved, PacBio-HiFi based *T. dactyloides* genomes [15]. These two genomes are of Tripsacum clones collected in Kansas (Td-KS) and Florida (Td-FL) representing northern and southern germplasm respectively. Within-and between-individual sequence divergence was estimated as in Stitzer et al (2025) for Andropogoneae tribe-wide syntenic genes. Briefly, *Paspalum vaginatum* genes were aligned to each assembly using AnchorWave (v1.2.3) [16], sequence was extracted for each gene region, and aligned using MAFFT v7.552 (parameters --genafpair --maxiterate 1000 --adjustdirection) [17]. We extracted codon positions based on the alignment and the *P. vaginatum* gene used as an anchor, and calculated synonymous substitutions between all pairwise combinations of gene copies using MSA2Dist [18].

The Illumina WGS paired end reads for founders, F1s, and bulks were aligned to a haplotype A of both the Td-KS and Td-FL genome assemblies using bwa-mem2 with default parameters. Read group information (sample, library, platform) was then added with gatk AddOrReplaceReadGroups [19,20], and alignments were sorted into BAM format using samtools [21]. For each founder, F1, and tolerant and susceptible bulk, sorted, read-grouped BAMs from the constituent individuals were merged per chromosome using samtools merge, and the resulting bulk BAMs were indexed with samtools index.

### Read QC

In order to verify pedigree and curate readsets, variants for each individual sequencing run of founders and F1s were identified using bcftools mpileup and bcftools call. To control for the varying sequencing depth across samples when calculating relatedness, all heterozygous sites for each sample in the VCF were randomly set to homozygous for one of the two alleles. Distance and normalized IBS between samples were then estimated using Tassel-5 [22], and non-parametric multidimensional scaling were performed using isoMDS from the MASS package [23,24]. Distance and IBS were used for manual curation of read-sets by checking that first-order relationships between samples met pedigree expectations.

### Alignment & Variant calling

**Figure 3:**
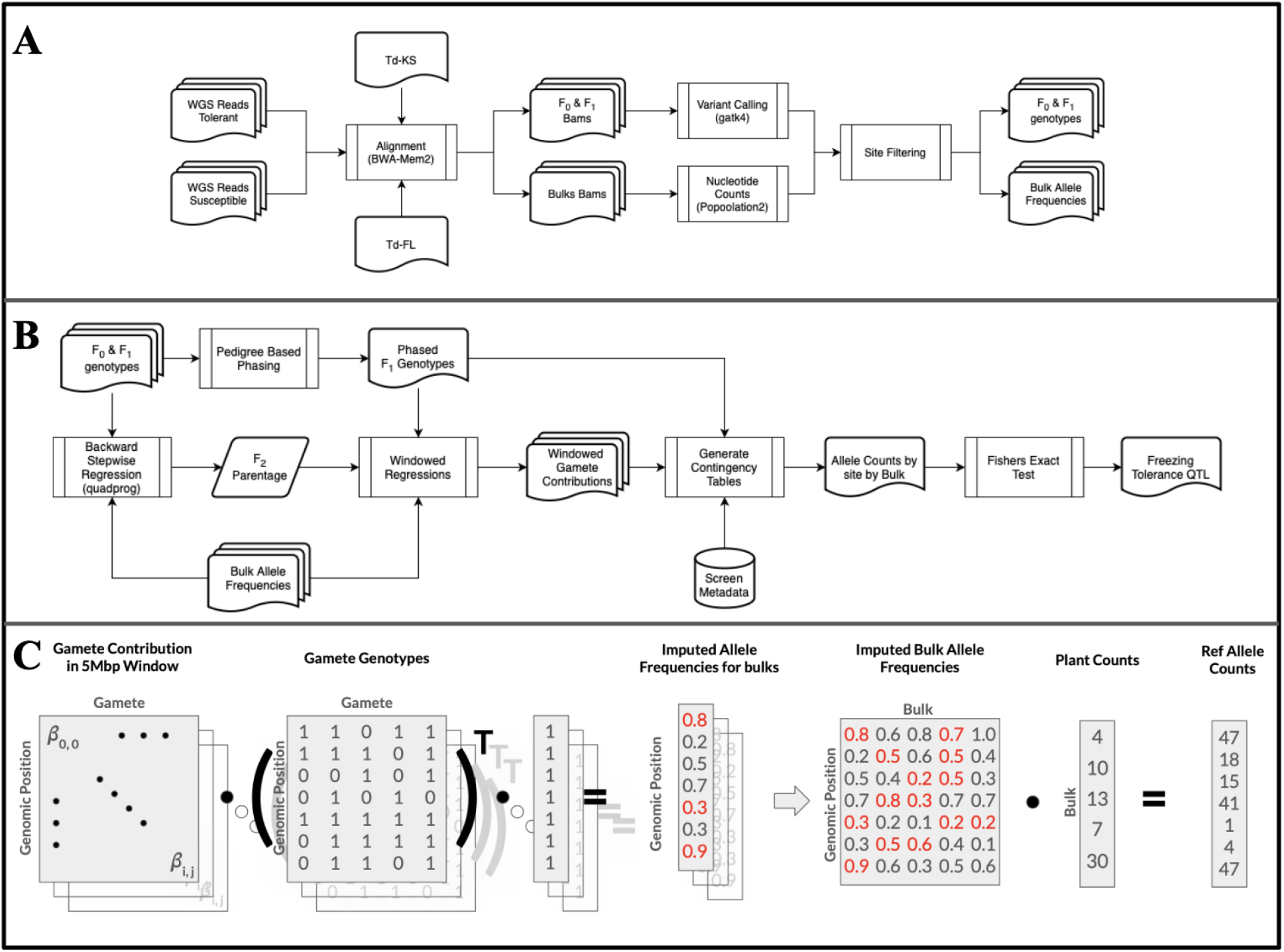
Overview of alignment and analysis pipeline going through the A) genotyping pipeline, B) BSA pipeline, and C) regression based allele frequency estimation and allele counting.

### Variant Calling of Founders and F1s

Variant calling and joint-genotyping was done using the GATK4 pipeline following the recommended best practices [20]. All samples were aligned and variants called relative to both the Td-KS and Td-FL reference genomes.

### Site Filtering of Founders and F1s

To avoid reference bias and select for quality SNPs, site filtering was done by checking that genotype calls against both references agree for each site. In order to test that genotype calls agree across reference genomes, positions from one genome needed to be translated to the coordinate system of the other genome. This was achieved by using AnchorWave to align the two genomes to one another and produce a multi-alignment format (MAF) file. This MAF file was then used to produce a chain file that allowed for conversion between coordinate systems. We then uplifted the genotype calls from Td-KS coordinates to Td-FL coordinates and subsequently tested sites for genotype call agreeance. Additionally, repetitive regions were filtered by using SEQbility [25] to produce a repeat mask in bed format. The identified non-repetitive genomic regions were intersected with the set of sites with genotype agreeance using bedtools.

To further refine our confident SNP set we looked at the distribution of site allele depth (AD) ratios in a set of 15 of our more deeply sequenced F1 individuals. If a site fell outside of the 5th and 95th percentile in 5 or more samples it was determined to be an untrustworthy site and removed from the dataset. The resulting high-confidence variant set became the foundation for the subsequent bulk segregant analysis (BSA).

### Variant Calling of Bulks

In order to overcome the challenge of variant calling within bulked samples that violate the ploidy assumption in most variant callers, a custom approach to genotyping was taken for the bulks. The merged and sorted BAMs were passed through samtools mpileup to produce the input for Popoolation2 [26]. The mpileup output was then provided to the mpileup2sync utility of Popoolation2 using the “fastq-type sanger” setting and minimum base quality of 20 to produce an allele depth table. This table was then used to calculate allele frequency for each bulk by using the depth of the reference allele over the total depth. We then rely on the previously identified high-confidence variant sites from the individually sequenced founders and F1s to filter sites in the bulks.

### Identification of Putative Temperature Adaptation Loci

Assuming the founder germplasms have been locally adapted to their original collection sites, genetic loci that have diverged among them indicate putative temperature-adaptation loci. We also noticed that *Tripsacum dactyloides subsp floridanum* was phenotypically distinct from *Tripsacum dactyloides subsp dactyloides* of southern origin collected from Florida. To further our understanding of these germplasm, we opted to subset our founders into three sub-groups (Sup Table 1), *Tripsacum dactyloides subsp dactyloides* of northern origin (TdN), *Tripsacum dactyloides subsp dactyloides* of southern origin (TdS; Florida and Texas origin), and *Tripsacum dactyloides subsp floridanum* (Tf; Florida everglades region).

To identify divergent regions among TdN, TdS, and Tf, we calculated Weir and Cockerham’s Fst for 5Kbp non-overlapping windows based on the biallelic SNPs using vcftools. These windowed estimates are then used to observe divergence trends across abiotic stress response genes.

### Regression-based gamete and allele frequency estimates

As a consequence of the open-pollinated nature of this experiment’s design, paired with the sparse sequencing of bulks, we are unable to accurately trace paternal contribution to the bulks, and we are unable to accurately estimate the allele frequency at any given site. To approximate these, we devised a two-step strategy leveraging the genotypes of the more deeply sequenced F1 and founder individuals.

### Whole Genome Backward Stepwise Regression

The first step is to estimate the parentage for each of the open-pollinated F2 bulks. We do this through a whole-genome backward stepwise regression.

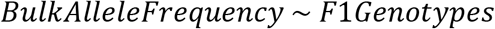

We regress the F1 genotypes onto the observed allele frequency of a given bulk while iteratively removing the least informative F1 based on the effect removal of that predictor has on the Bayesian information criterion (BIC). We designate the union of the top 3 predictors for both the tolerant and susceptible bulk of a family as the likely parentage. With the resulting parentage estimates, we further estimate the gametic proportion contributed by each parent for each bulk through a final genome-wide regression, but with some additional constraints [27]. We require: all regression coefficients (***β***) to be positive; the maternal parent’s regression coefficient to be at minimum 0.5; and the sum of all coefficients to be 1. This ensures that these regressions more accurately represent the biology of transmission genetics and assists in ease of comparison between families. We then also use the resulting regression coefficients to observe the spatial relationships of inter-crossing in the field. Using the estimated contribution frequencies of F1s to the bulks, we can begin to look for variation in contributions across the genome and determine putative freezing tolerance QTL by comparing relative contributions between bulks.

### F1 Phasing & Windowed Constrained Regressions

In order to improve our ability to resolve haplotypic contributions of the F1s, we first leverage the genotypic information of the population founders to phase the observed F1 genotypes to maternal and paternal founder gametes. Heterozygous calls were resolved by evaluating parental contributions. In cases where both parents were homozygous but disagreed, the F1 heterozygote was phased according to the parental configuration. When one parent was homozygous and the other heterozygous, the F1 haplotype was phased accordingly. If both parents were heterozygous, a default phased configuration was applied unless additional data justified a more precise assignment. Missing parental data were handled by imputing the most likely allele based on F1 genotype and available parental information. Sites with inconsistencies–such as heterozygous calls that contradicted parental contributions–were flagged and removed.

A regression model was applied independently for each genomic window (*i*), each family (*j*), and each bulk (*b*). In this model, the vector of bulk allele frequencies (*Y*_*ijb*_) within a given window was expressed as a linear combination of the gamete genotype matrix (*X*_*ijb*_) and a vector of regression coefficients (β_*ijb*_).

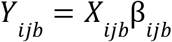

To perform the constrained regressions, the solve.QP() function from the quadprog package was used [27]. This function calculates the coefficients that minimize a quadratic objective function, representing the difference between the observed bulk allele frequencies and the predictions based on gamete genotypes, while ensuring that the coefficients satisfy specific constraints. In our analysis, we required that the coefficients for each family and bulk remain positive, sum to one, and that the sum of the maternal gamete coefficients is 0.5. By using solve.QP() with these constraints, we were able to derive coefficients that both adhered to known biological principles and accurately represented the gamete contribution frequencies. The difference of gamete contribution frequencies between susceptible and tolerant bulks evidences potential freezing-tolerance effect of a given window.

### Bulk Allele Frequency Imputation

Following the constrained windowed regressions, site-specific allele frequencies were imputed for each bulk. Using the regression-derived coefficients for each window, the contributions of individual gametes were applied to estimate the expected bulk allele frequencies. Specifically, for each genomic window, the observed gamete genotypes were multiplied by the corresponding regression coefficients to calculate the predicted allele frequencies for both tolerant and susceptible bulks. This provided a continuous, genome-wide estimate of allele frequencies and formed the basis for downstream statistical testing.

### Site-by-Site Significance Testing with Fisher’s Exact Test

To identify loci associated with freezing tolerance, Fisher’s exact tests were performed for each quality polymorphic site. First, a contingency table was created for every site, capturing the allele counts for both the tolerant and susceptible bulks. These tables reflected the summed contributions of the corresponding regression coefficients, weighted by the total number of alleles per bulk, as derived from plant count data. The fastfisher tool, a custom program, was then used to rapidly compute *p*-values for these contingency tables. The output from fastfisher provided a genome-wide set of *p*-values, allowing for the identification of putative freezing tolerance QTL.

### QTL Exploration

To guide interpretation of observed QTL, the genomic positions of known cold tolerance genes were determined by aligning their amino acid sequence, obtained from NCBI, against the Td-FL genome using miniprot [28].

To test for signals of abiotic stress adaptation, conserved stress response gene sets were defined for cold, drought, heat, and water logging. These genes were identified to consistently respond to stress across 41 expression studies of Poaceae species (8-11 studies per stress, we require significant responses in at least three studies per stress to be called as consistently responsive) (Hsu et al., in prep). Tests for significant deviations in divergence between stress-responsive and respective non-responsive genes were conducted through a *t*-test on 5kbp windowed Fst estimates overlapping for each set. Genes were assigned to a window based on their start coordinate.

## Results

### Germplasm genotyping

To avoid potential reference bias in genotyping, we included the genome assemblies of two distinct *Tripsacum dactyloides subsp. dactyloides* individual plants as the references. Td-KS was collected in Kansas and is representative of our northern population. Td-FL was collected in Florida and is representative of our southern population. Both genomes are highly contiguous, haplotype resolved, and have genes annotated. Comparing the references to one another, genome wide sequence divergence was estimated to be 0.017 with an average Ks of 0.0092 (Sup Figure 1). The synonymous substitution rate (Ks) between allelic gene copies for Td-KS and Td-FL is ∼0.0136 and 0.0045, respectively.

In total, we sequenced and genotyped 38 diverse founder accessions and 38 F1s using WGS short-read technology (Sup Table 5: mean sequencing coverage for each founder and F1). We separately mapped raw sequencing reads to both genomes and called ∼208M and ∼220M SNPs against Td-FL and Td-KS respectively. After filtering we only considered the ∼54M SNPs that have consistent genotype calls against both references, located outside of repeated regions and exhibit reasonable allele depth ratios across samples for subsequent analyses giving us a final density of ∼18 SNPs per 1 Kbp.

To identify genetic loci associated with freezing tolerance through bulk segregant analysis, 22 F2 families were put through freezing screens of which approximately 15-20% of the top and bottom performing individuals were selected for bulk-sequencing. Allele frequencies were calculated for each bulk on the ∼54M high-confidence sites.

### Genomic diversity of Tripsacum germplasm

To identify genetic relatedness, we performed an MDS analysis on the genotypes of these accessions. We observed three distinct clusters among the founder accessions (Fig 4). The three accessions of *T. dactyloides subsp. floridanum* are clustered. Closest to the subspecies *floridanum* cluster are a number of *T. dactyloides subsp. dactyloides* accessions that are generally distributed across the southern Coastal Plains. Distant from the first two clusters is a third cluster of *T. dactyloides subsp. dactyloides* that are found across the Great Plains.

**Figure 4:**
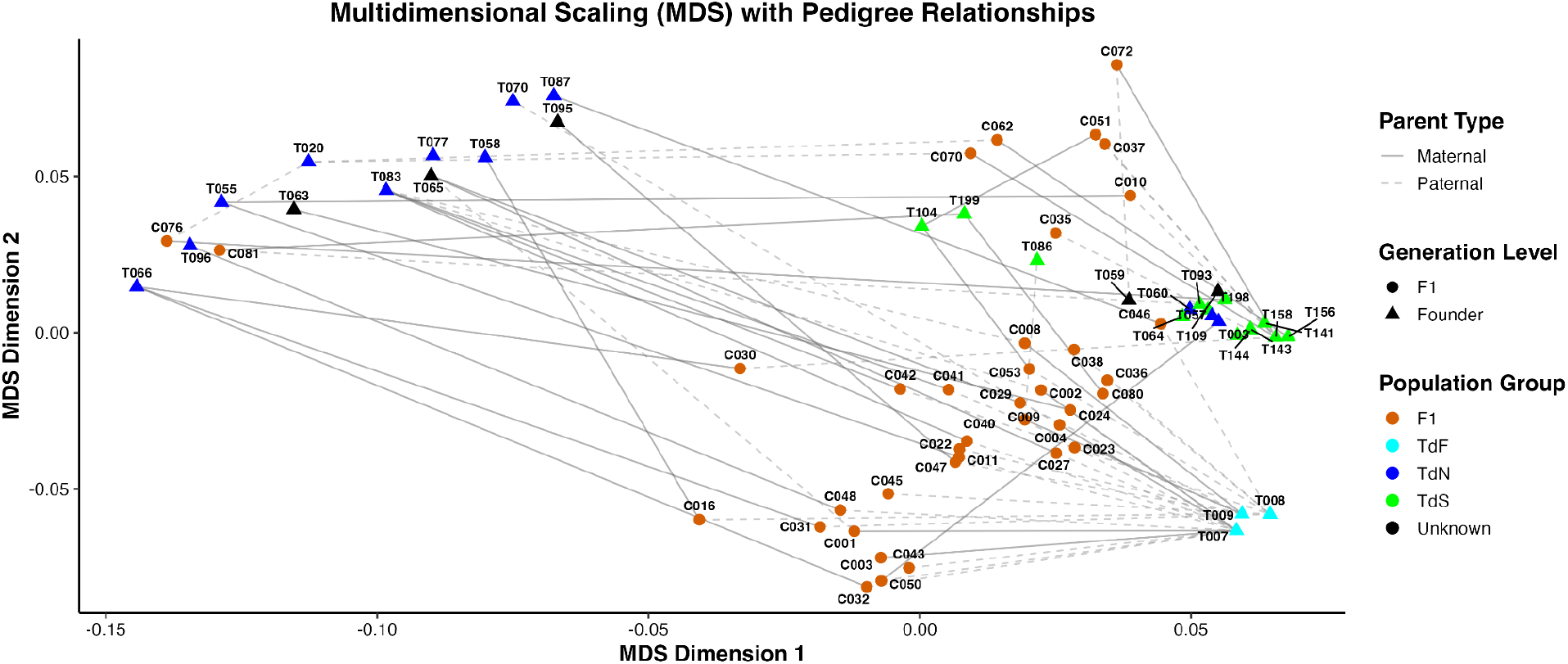
Non-parametric MDS generated by the isoMDS function from the MASS R package with distance matrix produced by Tassel-5. Lines connect F1 to their respective parents based on known pedigree information.

### Genome-wide divergence between northern and southern accessions

As the MDS analysis shows, latitude is a key driver behind the genomic divergence of the studied Tripsacum accessions. We further investigated the divergence among the sub-groups in these accessions through pairwise Fst estimates. All three groups exhibited relatively high differentiation (Fst of 0.25 to 0.30), however, it is notable that the Southern versus floridanum comparison exhibited higher variance with more genomic regions close to complete differentiation (Fig 5A). We observe a genome wide divergence between TdN and TdS (Weir and Cockerham weighted Fst estimate of 0.25). This genome wide divergence is more so pronounced when comparing the two clusters of Td to Tf (Weir and Cockerham weighted Fst estimate of 0.30 and 0.27 respectively; Fig 5A). When plotting weighted Fst values in 5kbp windows across the genome we observe distinct linkage patterns for each pairwise comparison (Fig 5C).

**Figure 5.**
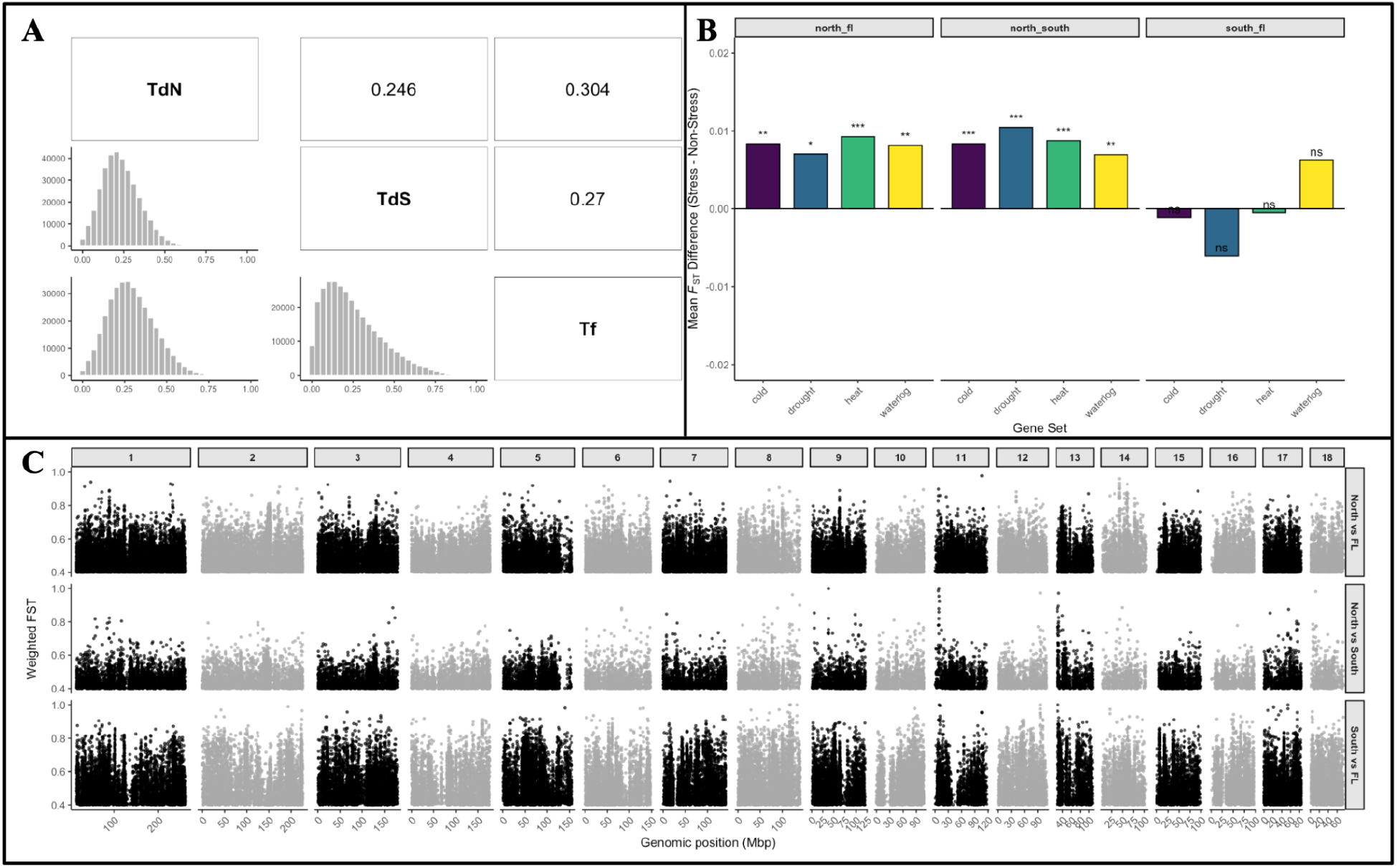
Genomic divergence and stress-responsive differentiation among Tripsacum groups. (A) Pairwise Weir-Cockerham Fst shows strong divergence among the three groups, with higher variance and more near-fixed regions in the Southern-floridanum comparison. (B) Mean windowed Fst is elevated for genes responsive to cold, drought, heat, and waterlogging when comparing Southern to Northern groups, but not when comparing Southern to *floridanum*. (C) Genome-wide 5-kb Fst windows reveal distinct divergence patterns for each pairwise comparison.

To test for signals of abiotic stress adaptation, we use sets of genes identified to consistently respond to a number of stresses across the Poaceae including cold (2549 genes), drought (1524 genes), heat (3575 genes), and water logging (3064 genes). Using a t-test and the windowed Fst estimates, we saw significant differences in divergence level between responsive genes and non-responsive genes. For both the southern groups against the northern subgroup, we observe a statistically significant and positive difference in mean Fst for all of the sets of stress response genes relative to all other genes. Conversely, between the Southern and floridanum groups, we observe no significant difference in Fst for any of the stress responsive gene sets (Fig 5B).

### Imputation of parental gametes

Based on a backwards regression approach, we identified the major contributing F1 parents in each of the 22 F2 families (A family is composed of the tolerant and susceptible bulks derived from a single maternal F1 source). Estimates of contribution frequency show the maternal F1 contributes the most in nearly all families and the following contributions usually come from F1s which are in adjacent rows in the field (Sup Fig 1: network and field).

### Segregation of freezing tolerance in F2

In order to observe segregation across the genome, the genotypes of the inferred contributing F1s are phased to the founder contributed gametes and used in 5Mbp windowed constrained regressions to reconstruct the segregating gamete frequencies and recombination patterns in each family (Fig 6).

**Figure 6.**
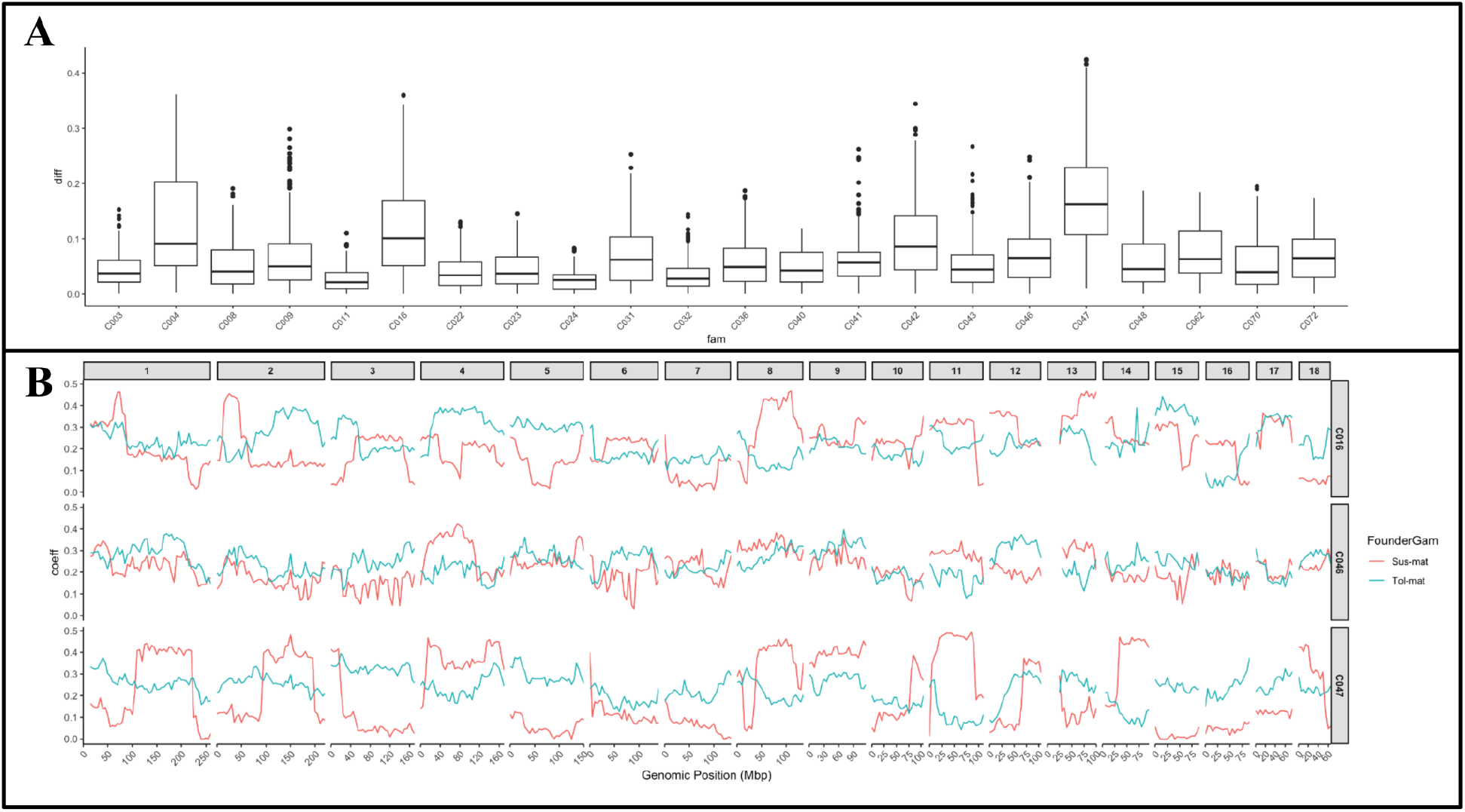
Maternal gamete segregation and recombination patterns across families. (A) Barplots show the maternal segregation index for all 5-Mb windows in each family. (B) Lineplots show maternal gamete segregation across all chromosomes for three families, with red tracing susceptible-bulk frequencies and blue tracing tolerant-bulk frequencies. Families differ in genome-wide segregation magnitude, in localized reductions in recombination, and in regions where high maternal IBD obscures segregation signals.

Although differences in segregation are observed, resolution of these patterns vary substantially by family and even by chromosome. This is most easily observed in the segregation frequency differential of the maternal gametes between bulks. We observe relatively elevated rates of genome-wide genetic segregation in some families (i.e. C047, C016, and C004), a number of families that seem to have depressed rates of recombination (i.e. C047 chr15), and others that show high IBD in the maternal F1 genotype (i.e. C011, C024) consequently making recombination and segregation patterns unidentifiable.

### Joint bulk segregant analysis identified freezing tolerance-associated QTLs

Since gamete estimations are done in equal 5Mbp windows across families, we can identify regions of the genome that consistently show high segregation between bulks by investigating the relative difference in contribution frequency of the maternal F1 gametes in each particular 5Mbp window (Fig 7A; Sup Table 6). We observe elevated rates of segregation around a few classically known cold responsive genes including the Cold Binding Factors (CBFs). We also tested for enriched segregation rates of genes that consistently respond to environmental factors across the Poaceae (Hsu et al., in prep). We observed a marginal positive shift in cold response genes and water-logging response genes when compared to all genes, but no shift in drought and heat response genes when compared to all genes.

**Figure 7.**
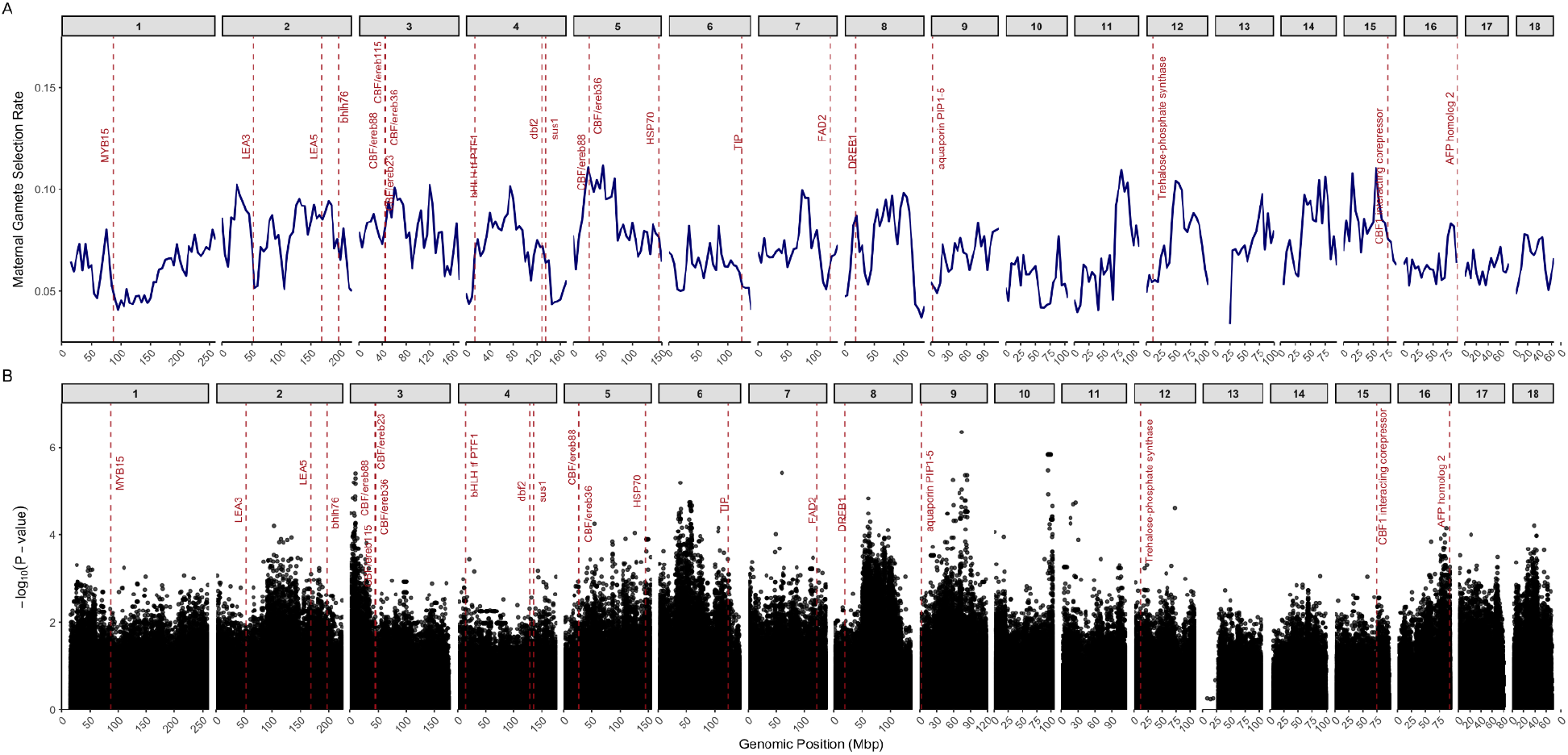
Genome-wide segregation peaks and allele-frequency divergence associated with freezing tolerance. (A) Maternal gamete segregation rates in 5-Mb windows highlight genomic regions that consistently show elevated segregation between bulks, including intervals near known freezing-tolerance genes such as the CBF cluster. (B) Site-specific allele-frequency tests using imputed F2 bulk frequencies identify 9 significant QTL (p < 1×10^−5^) for freezing-tolerance segregation. Most QTL span 1-4 Mb and show consistent allele-frequency divergence across families.

These reconstructed gamete frequencies enabled effective imputation of the allele frequencies in the F_2_ bulks (Fig 2C) which allows for site-specific allele segregation testing. With the imputed allele frequency table, we performed a joint site-by-site fishers-exact test which tested for divergence in allele frequency between the tolerant and susceptible bulks. At the significance threshold of p < 1e-5, we identified 9 QTL significantly associated with the segregating of freezing tolerance (Fig 7B; Sup Table 7). On average, we observed clear frequency divergence of the significant QTLs across the tested families. Independent recombination events in different families help increase the mapping resolution while most identified QTLs span 1-4 Mbps in the genome, preventing further candidate gene nomination.

## Discussion

Our study identified multiple genomic regions associated with freezing tolerance in Tripsacum dactyloides. Joint maternal gamete segregation highlighted numerous genomic regions with differential segregation (Fig 7A), and joint allele frequency tests identified 9 quantitative trait loci (QTLs) significantly segregating between freezing-tolerant and susceptible seedlings (Fig 7B). These QTL provide candidate genetic resources for enhancing maize resilience to frost. However, substantial genetic divergence and limited recombination within our wild-derived mapping population reduced mapping resolution. These limitations underscore challenges in mapping adaptive traits in undomesticated, heterozygous species.

### Characterization of Tripsacum Subpopulations

Through genetic analysis of Tripsacum dactyloides germplasm, we identified three genetically distinct populations that seem to generally center around three geographic regions: (1) the Great Plains (central grasslands), (2) the Coastal Plains (Gulf and Atlantic coastal grasslands), and (3) southern Florida’s coastal plains with Tripsacum dactyloides subsp. floridanum, an endemic subspecies restricted to. These contemporary populations may be derived from ancestral populations isolated likely originated from two ancestral refugial populations isolated around the Last Glacial Maximum (∼18 kya [29]): a western lineage distributed across the Gulf and Atlantic Coastal grasslands, and an eastern lineage associated with the Florida Peninsula refugium, potentially separated by the Mississippi River discontinuity [14,30]. Subsequent admixture events between these ancestral lineages presumably led to the formation of the present day Great Plains and Coastal Plains populations while, in contrast, *T. dactyloides subsp. floridanum* underwent geographic isolation in southern Florida, resulting in ecological specialization to arid, sandy coastal habitats. Further resolution of the migratory history and adaptive evolution of these ecotypes requires increased sampling density across additional key ecoregions, notably the Southeastern USA Plains (Ecoregion 8.3), Ozark and Ouachita-Appalachian Forests (Ecoregion 8.4), and Mississippi Alluvial and Southeast USA Coastal Plains (Ecoregion 8.5).

### Limitations to High Resolution Gene Mapping

We expected that each cross between northern and southern populations would result in independent recombination events and consequently, comparing QTL across these recombination events would improve our resolution and capacity to identify candidate genes. Instead, we observed independent recombination patterns in some crosses, but others retained nearly complete haplotype identity when compared to the haplotype contributed by their respective founder (Fig 6B). Given our data, it is unclear whether this outcome results from strong selection bias toward one parental genotype or reflects biological barriers, such as recombination suppression. Because our crosses often are made between T. dactyloides subspecies (subsp. dactyloides × subsp. floridanum), these patterns may reflect nascent reproductive isolation mechanisms between these lineages.

Another unexpected observation comes from the limited concordance between loci identified through maternal gamete segregation (Fig 7A) and association tests (Fig 7B). This discrepancy may result from the extensive genetic variation within the mapping population as depicted in our Fst results (Fig 5). Multiple alleles with equivalent phenotypic effects might segregate simultaneously, reducing our power to pinpoint specific causal variants, yet allowing detection of broader segregation patterns of a particular locus. Indeed, classical cold-response genes prominently appear in maternal gamete segregation indices, despite absence from significant allele segregation results.

We observe that this population does segregate phenotypically and genotypically for freezing tolerance and could be a key source for developing freezing tolerant maize, but the functional variation that confers freezing tolerance across Tripsacum accessions may differ. It is possible that Tripsacum by-and-large is freezing tolerant and what we are observing are a number of independent loss-of-function events causing susceptibility. While this would make it unlikely that putative alleles identified in Tripsacum would be the key to creating freezing tolerant maize, it may help narrow down to particular molecular pathways. This population thus serves as a valuable genetic resource, yet further studies must both identify putative Tripsacum-derived freezing tolerance alleles and confirm whether they confer similar phenotypes when transferred into maize (Zea mays).

Characterization of the Tripsacum subpopulations could be better resolved with more dense and distributed genotyping of existing Tripsacum diversity, whether this be whole genome assemblies or skim sequencing. Improvements toward resolving existing Tripsacum subpopulations would benefit both the creation of targeted mapping populations, and potentially improve Fst resolution. Additionally, while the QTL identified in this study remains too large to identify putative causal mutations, we do observe differential constraint and segregation of conserved stress responsive genes suggesting these QTL could be further resolved to identify putative mutations. Reasonable approaches to do this include expression GWAS, proteomics, metabolomics along with functional characterization of mutations contributing to differential expression of significant independent response genes. Once these putative causal mutations or response pathways are identified, work to explore the functionality of these pathways can be done to determine genetic engineering based approaches to improve maize hardiness to freezing conditions.

## Acknowledgments

We thank Peter L. Putriment for his contributions in designing the freezing screen protocol.

This work was funded by: the NSF PanAnd Grant Award #1822330; the NSF National Research Traineeship in Digital Plant Science Award #1922551; the USDA-ARS.

This work utilized both Cornell BioHPC and SCINet for compute resources.

